# Synthetic lethality-based prediction of anti-SARS-CoV-2 targets

**DOI:** 10.1101/2021.09.14.460408

**Authors:** Lipika R. Pal, Kuoyuan Cheng, Nishanth Ulhas Nair, Laura Martin-Sancho, Sanju Sinha, Yuan Pu, Laura Riva, Xin Yin, Fiorella Schischlik, Joo Sang Lee, Sumit K. Chanda, Eytan Ruppin

## Abstract

Novel strategies are needed to identify drug targets and treatments for the COVID-19 pandemic. The altered gene expression of virus-infected host cells provides an opportunity to specifically inhibit viral propagation via targeting the synthetic lethal (SL) partners of such altered host genes. Pursuing this antiviral strategy, here we comprehensively analyzed multiple *in vitro* and *in vivo* bulk and single-cell RNA-sequencing datasets of SARS-CoV-2 infection to predict clinically relevant candidate antiviral targets that are SL with altered host genes. The predicted SL-based targets are highly enriched for infected cell inhibiting genes reported in four SARS-CoV-2 CRISPR-Cas9 genome-wide genetic screens. Integrating our predictions with the results of these screens, we further selected a focused subset of 26 genes that we experimentally tested in a targeted siRNA screen using human Caco-2 cells. Notably, as predicted, knocking down these targets reduced viral replication and cell viability only under the infected condition without harming non-infected cells. Our results are made publicly available, to facilitate their in vivo testing and further validation.

## Introduction

The coronavirus disease 2019 (COVID-19) pandemic caused by the novel coronavirus SARS-CoV-2 together with its emerging new variants in 2021 has resulted in hundreds of millions of infected people with millions of deaths worldwide (WHO Coronavirus Disease (COVID-19) Dashboard, 2021). Up to this date, there are several thousand registered clinical trials of anti-COVID-19 therapies worldwide (International Clinical Trials Registry Platform (ICTRP), 2021), with remdesivir being the only drug approved by the United States Food and Drug Administration (FDA) and the drug regulatory agencies of several other countries (Beigel et al. 2020). However, the clinical benefit of remdesivir in most COVID-19 patients is still modest (Pan et al. 2020). A total of 11 therapies, including virus-neutralizing monoclonal antibodies bamlanivimab and etesevimab (Mahase, 2020), sotrovimab, the combination of casirivimab and imdevimab, interleukin-6 receptor inhibitor Tocilizumab and Janus kinase (JAK) inhibitor baricitinib (in combination with remdesivir) have obtained Emergency Use Authorization (EUA) from the U.S. FDA (U.S. Food and Drug Administration, 2021). Dexamethasone and other corticosteroids have been recommended for the control of COVID-19-related symptoms (Ledford, 2020). Numerous preclinical studies have conducted anti-SARS-CoV-2 drug repurposing as well as genetic screening of varied scales (for example, Riva et al. 2020; Wei et al. 2020; Danoliski et al. 2020), supplemented by other studies providing computational target predictions (for example, Zhou et al. 2020; Tilocca et al. 2020; Zhou et al. 2020a; Cheng et al. 2021a). Two mRNA vaccines from Pfizer-BioNTech and Moderna have been approved by the U.S. FDA (U.S. Food and Drug Administration, 2021). Despite all these efforts, there is an urgent unmet need to identify additional anti-SARS-CoV-2 targets and drugs, to resolve the COVID-19 crisis (Jomah et al. 2020).

Numerous recent studies have contributed to the understanding of the biology of SARS-CoV-2 infection (for example, Paules & Fauci, 2021; V’kovski et al. 2020; Bojkova et al. 2020; Forster et al. 2020). A lot of interest has been focused on targeting *ACE2* and *TMPRSS2*, the virus entry receptors (Muralidhar et al. 2021; Ragia & Manolopoulos, 2020) and the search for drugs targeting these genes is underway (Lin et al. 2021; Hoffman et al. 2020). However, the direct targeting of the viral life cycle to inhibit its proliferation remains challenging. Alternatively, as viruses are known to alter the expression and activity of many host genes upon infection, it has been suggested that one may selectively target the infected cells by targeting the genes that are synthetic lethal (SL, including synthetic dosage lethal, SDL) with the virus-altered genes (i.e. their “SL or SDL partners”) (Mast et al. 2020). SL denotes a genetic interaction between two genes, where the inactivation of both genes, but not either gene alone, leads to decreased cell viability (illustrated in **Figure 1a**), so by targeting the SL partners of a gene that is markedly down-regulated specifically in infected cells only, one may selectively place the infected cells under stress without harming non-infected cells, thus diminishing the capacity of the infected host cells to support viral proliferation and inhibiting viral propagation (Mast et al. 2020; illustrated in **Figure 1a**). A variant of SL, called synthetic dosage lethality (SDL), is another type of genetic interaction, where the overexpression of one gene and the inactivation of a second gene reduces the cell viability (Kroll et al. 1996). Here we use SL as a general term referring to both SL and SDL. A proof-of-concept study has demonstrated the value of this strategy for the inhibition of poliovirus (Navare et al. 2020). We aim to systematically identify SL based targets for inhibiting SARS-CoV-2 infection.

**Figure 1.**
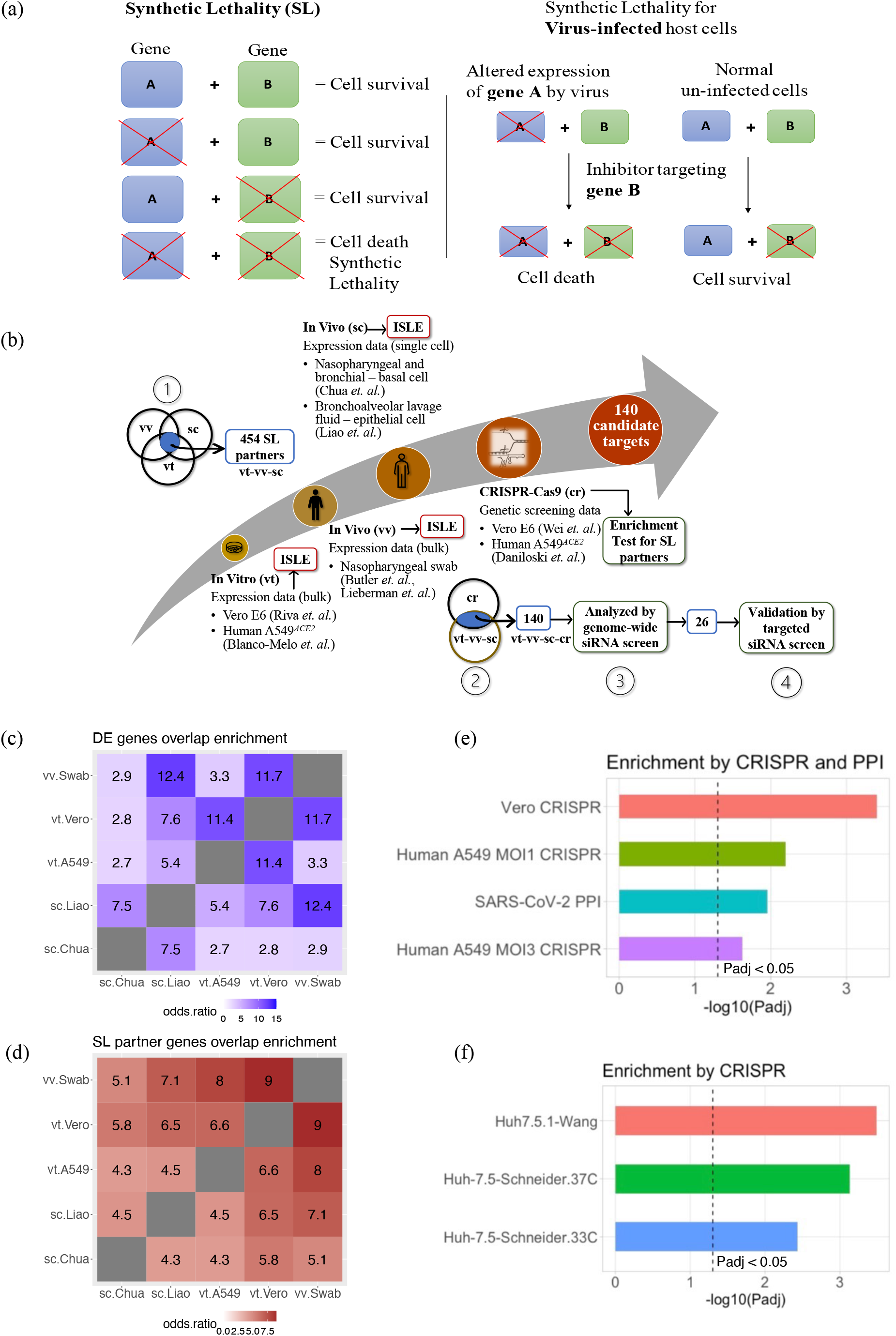
**(a)** An illustration of the concept of synthetic lethality (SL, left-hand side) and its application to the context of anti-viral infection (right-hand side). **(b)** An illustration of the overall workflow we used to identify SL-based candidate targets for anti-SARS-Cov-2 by integrating different in vitro (vt) and in vivo (vv), including single-cell (sc) datasets. Highlighted modules are: (1) Intersection of ISLE-predicted SL/SDL partners of the SARS-Cov-2-induced differentially expressed (DE) genes in all the denoted vt, vv and sc datasets yielded 454 genes (vt-vv-sc). (2) These 454 vt-vv-sc genes are shown to be enriched for strong hits from two different CRISPR-Cas9 screens (cr) in the SARS-CoV-2 infection setting (see main text for details), and the intersection with the hits from the CRISPR screens yielded 140 final candidate SL-based anti-SARS-CoV-2 targets (vt-vv-sc-cr). (3) These 140 candidates are further filtered based on their enrichment for genes whose knockdowns decrease cell viability in a genome-wide siRNA screen we performed earlier. (4) We further selected a subset of 26 targets from genome wide siRNA screen and validated experimentally via a small-scale targeted siRNA screen. **(c)** A heatmap showing the extents of overlaps between the differentially expressed (DE) genes in SARS-CoV-2 infected samples vs non-infected controls identified from different datasets, as measured by the odds ratio of enrichment between each pair of datasets, which is encoded by the color and also labeled in the cells within the heatmap. The dataset labels are as follows: vt.Vero (Vero E6 cell samples from Riva et al. 2020), vt.A549 (A549^ACE2^ cell samples from Blanco-Melo et al. 2020), vv.Swab (COVID-19 patient nasopharyngeal swab samples from Butler et al. 2020 and Lieberman et al. 2020), sc.Chua (single-cell data of nasopharyngeal swab and bronchial samples from Chua et al. 2020) and sc.Liao (single-cell data of bronchoalveolar lavage fluid samples from Liao et al. 2020). **(d)** A heatmap showing the extents of overlaps between the SL/SDL partner genes identified with ISLE based on the different datasets, with the same dataset labels as in (c). **(e)** The negative log10-transformed Benjamini-Hochberg-adjusted P values (Padj) from Fisher’s exact tests (X-axis) for the enrichment between the 454 consensus ISLE-identified SL-based candidate targets and different validation gene sets, including: genes with strong negative log fold-changes identified in the CRISPR-Cas9 screen in SARS-CoV-2-infected Vero E6 cells from Wei et al. 2020 (vero CRISPR), such genes in another CRISPR-Cas9 screen in SARS-CoV-2-infected A549^ACE2^ cells (Daniloski et al. 2020) at two different multiplicities of infection (Human A549 MOI1 CRISPR and Human A549 MOI3 CRISPR, respectively), and human genes interacting with SARS-CoV-2 proteins identified in Gordon et al. 2020 and Stukalov et al. 2020 (SARS-CoV-2 PPI). The black dotted line corresponding to the cutoff of adjusted P<0.05. **(f)** The negative log10-transformed Benjamini-Hochberg-adjusted P values (Padj) (X-axis) resulting from a GSEA enrichment analysis between the 140 candidates and genes with strong negative log fold-changes identified in the CRISPR-Cas9 screen in SARS-CoV-2-infected human Huh-7.5.1 hepatoma cells from Wang et al. 2021 (Huh7.5.1-Wang), and genes in another CRISPR-Cas9 screens in SARS-CoV-2-infected human Huh-7.5 hepatoma cells (Schneider et al. 2021), at 37° C and 33° C (Huh-7.5-Schneider.37C and Huh-7.5-Schneider.33C, corresponding to two physiologically relevant temperatures of the lower and upper airways, respectively). The black dotted line corresponding to the cutoff of adjusted P<0.05. These two CRISPR-Cas9 screens were not used to generate the list of 140 targets.

Previously, we have developed a computational pipeline named ISLE (identification of clinically relevant synthetic lethality) to identify clinically relevant SL genetic interactions by mining large-scale genomic and transcriptomic data across human cancers (Lee et al. 2018). Applying ISLE, we have previously successfully identified a Gαq-driver mutation as marker for FAK inhibitor in uveal melanoma (Feng et al. 2019) and a synergistic combination of asparaginase and MAPK inhibitors for treating melanoma and pancreatic cancer (Pathria et al. 2019). Although the ISLE algorithm was originally designed to predict SL in cancers, we have recently found that some SL gene pairs predicted by ISLE are also functional in non-cancerous tissues (Cheng et al. 2021b), reflecting the conservation of the wiring of molecular networks of human cells more generally. Therefore, we reasoned that ISLE can also be valuable in predicting SL gene pairs for non-cancer contexts. Adopting this conjecture, here we applied ISLE to a large cohort of both *in vitro* and *in vivo* SARS-CoV-2 RNA-sequencing datasets to identify SL-based anti-SARS-CoV-2 targets. We then set to combine these predictions with the results of large-scale SARS-CoV-2 *in vitro* genome-wide CRISPR and siRNA screens. Focusing on a selected list of 26 predicted targets, we tested them experimentally in a small-scale targeted siRNA screen and demonstrated their efficacy in inhibiting SARS-CoV-2 replication. As these targets are supported by analyzing *in vivo* data, these are more likely to be of translational relevance.

## Results

### Overview of our strategy for SL-based anti-SARS-CoV-2 target prediction

Our analysis proceeds in four steps: **(A)** First, we analyzed multiple published host cell gene expression datasets (**Figure 1b**) involving *in vitro* and *in vivo* human samples of SARS-CoV-2 infection to identify the strongest significantly differentially expressed (DE) genes in response to viral infection. **(B)** Second, applying the ISLE pipeline, we identified the most robust clinically relevant SL partner of these DE genes in human cells, as potential SL-based anti-SARS-CoV-2 targets. We show that these 454 predicted targets (**Figure 1b**, module 1) are enriched amongst genes whose knockout <u>strongly reduced cell viability</u> in several SARS-CoV-2 infection CRISPR-Cas9 genetic screens. **(C)** We further filtered and prioritized a smaller list of 140 predicted targets based on support by several CRISPR screens (**Figure 1b**, module 2), and a high throughput genome-wide siRNA screen conducted and analyzed in our lab (**Figure 1b**, module 3). **(D)** Finally, we employed a network-based simulated annealing algorithm to further prioritize 26 highly ranked targets (**Figure 1b**, module 4), and conducted a small-scale targeted siRNA screen to experimentally test these targets in greater depth. A stepwise detailed description of the analysis performed, and the results obtained in each step follows.

### (A) Identification of host genes differentially expressed after SARS-CoV-2 infection

To identify the host cell genes whose expression is altered upon SARS-CoV-2 infection, we collected and analyzed multiple transcriptomic datasets of SARS-CoV-2 infection (**Figure 1b**). These include bulk RNA-seq data of *in vitro* samples from Vero E6 cells (Riva et al. 2020) and human A549 cells with exogenous *ACE2* expression (A549^*ACE2*^) (Blanco-Melo et al. 2020), as well as *in vivo* nasopharyngeal swab samples from COVID-19 patients (Butler et al. 2020, Lieberman et al. 2020). For *in vivo* analysis we additionally used single-cell RNA-seq data of nasopharyngeal and bronchial samples (Chua et al. 2020) and bronchoalveolar lavage fluid (BALF) samples (Liao et al. 2020). For each of these datasets, we identified the significantly differentially expressed (DE) genes in the SARS-CoV-2-infected samples vs matched virus-negative controls (FDR<0.05 with additional dataset-specific criteria, see Methods). For the single-cell datasets, we performed the DE analysis only in the epithelial cells, as these represent a primary target for SARS-CoV-2 infection, thus being our focus for SL-based antiviral strategy. Comparing the identified DE genes across datasets with Fisher’s exact tests (Methods), we find varied but overall higher-than-random pairwise similarities (i.e. odds ratio >1, **Figure 1c**). The lowest odds ratio (2.7, adjusted P=2.66E-03) is found between the *in vitro* A549^*ACE2*^ cell dataset and the *in vivo* single-cell RNA-seq samples (the Chua et al. dataset), and the highest odds ratio (12.4, adjusted P=1.33E-38) is seen between the in vivo single-cell (the Liao et al. dataset) and in vivo bulk swab datasets. Reassuringly, the DE genes identified from the in vivo bulk and single-cell RNA-seq datasets have reasonably good overlaps (**Figure 1c**). We performed pathway enrichment analysis of the DE genes from each dataset (Methods; **Table S1**) and noticed that although these genes are different across datasets, they are enriched for functionally similar pathways (e.g., platelet aggregation plaque formation pathway in human A549^*ACE2*^ dataset and platelet activation signaling and aggregation in *in vivo* single-cell Chua et al. dataset; **Table S1**).

### (B) Identification of the SL partners of SARS-CoV-2-induced DE genes

Next, we predicted the SL partners of the identified DE genes, as they constitute potential antiviral targets that can selectively impair the function of infected host cells and thus limit their capacity to support viral proliferation (list of SL pairs of each dataset are given in **Table S2**). Specifically, we applied the ISLE algorithm (Lee et al. 2018) to predict the SL partners of the genes significantly down-regulated by SARS-CoV-2 infection in the datasets described above. For the up-regulated genes, we analogously predicted their SDL partners with a modified version of ISLE (Methods). Interestingly, the SL partners identified across datasets with FDR cutoff 0.1 exhibit overall higher similarities than those similarities observed among the DE genes, with a minimum pairwise odds ratio of 4.3 (adjusted P=1.48E-118; **Figure 1d**). In particular, the overlap in the SL partner genes between the *in vitro* A549^*ACE2*^ cell dataset and the *in vivo* single-cell RNA-seq samples (in Chua et al. dataset) is higher than the overlap observed between their corresponding DE genes (SL odds ratio of 4.3 compared to corresponding DE odds ratio of 2.7). This suggests that the DE genes identified across the datasets may be functionally similar, resulting in higher chances of forming SL interactions with the same genes, which is indeed supported by the pathway enrichment results of the DE genes as described above (**Table S1**). Reassured by the overall similarity of the SL partner genes across datasets, we then selected a subset of 454 SL/SDL partner genes that is common to all in vitro and in vivo datasets (**Table S2F**, see details in Methods; illustrated in **Figure 1b**, module 1), which we focus on further analysis.

### (C) Initial screening and filtering of predicted SL-based targets with published CRISPR-Cas9 genetic screens

To further filter the 454 predicted SL-based targets, we collected data from two published CRISPR-Cas9 genetic screens that aim to identify host genes modulating SARS-CoV-2 replication. These include a screen in the Vero E6 cell line (Wei et al. 2020) and another in the human A549^*ACE2*^ cell line at two different multiplicities of infections (MOIs; Daniloski et al. 2020). We note that these studies seek to identify targets whose knockout promotes cell survival by reducing the virus-induced cytopathic effects, while our strategy aims to identify genes whose knockout specifically reduces the viability of infected cells. Thus, we expect that our predictions would be enriched in genes whose knockout results in strong cell dropouts in these genetic screens. Indeed, we find that our 454 predicted SL-based target genes are significantly enriched for cell viability-reducing genes (adjusted P values from Fisher’s exact tests range from 2.38E-02 to 4.00E-04, **Figure 1e**; Methods). The 454 SL-based targets are also enriched for host proteins that interact with SARS-CoV-2 viral proteins (Fisher’s exact test adjusted P=1.10E-02, **Figure 1e**; data of virus-interacting proteins is from Gordon et al. 2020 and Stukalov et al. 2020), indicating that the predicted targets are closely relevant to SARS-CoV-2 biology and infection. Among the 454 predicted targets, 140 genes are significantly strong cell viability-reducing genes (Wei et al. 2020, P=1E-05; Daniloski et al. 2020, P=0.002 for MOI1 and P=0.02 for MOI3) when depleted in both CRISPR-Cas9 screen datasets (illustrated in **Figure 1b**, module 2; **Figure 1e**). These genes are also significantly enriched amongst the genes whose knockout reduced cell viability in two other recently published genome-wide CRISPR-Cas9 knockout screens in human hepatoma cells (Wang et al. 2021, P=1.09E-04; and Schneider et al. 2021; P=3.73E-03 for the screen at 33° C and P=4.95E-04 for the screen at 37° C; **Figure 1f**). We thus focused our further analysis on these 140 candidate targets as listed in **Table S3**, ranked by a weighted DE score (WDS) that considers both the number of DE genes genetically interacting with each SL partner and the DE fold change (Methods).

Among those 140 targets, those that are inhibited by existing drugs are listed in **Figure 2a (Table S3B)**, together with these drugs’ mechanisms of action, as potential candidates for drug repurposing. Among these inhibitory drugs, warfarin and dicoumarol, targeting *VKORC1* are FDA approved. There is one clinical trial for Covid-19 patients on chronic treatment with anticoagulant warfarin vs who are not receiving (NCT04518735). Other experimental drugs are undergoing clinical trials either for cancer or other diseases. Notably, quercetin, targeting *CEBPB* is specifically undergoing 12 clinical trials (such as NCT04844658, NCT04578158, NCT04377789) for Covid-19 patients. A pathway enrichment analysis of these 140 SL-based targets (Methods, **Table S3E**) shows enrichment in pathways including cell cycle processes, stress responses, DNA break repair processes, RNA metabolic processes and RNA polymerase II transcriptions (**Figure 2b**). The DE genes are enriched in pathways including complement cascade, ion channel transport, innate immune processes, cytokine signaling, interferon signaling and neutrophil degranulation (**Figure 2c, Table S3F**). A pairwise pathway enrichment analysis (Methods; **Figure 2d**) shows that many SL interactions include cell-cell communications, innate immune system, post translational protein modification (i.e., columns with high numbers of significant P values in **Figure 2d, Table S3G**). The pathway combination involving the largest number of gene pairs is between the innate immune system (DE genes) and the cell cycle (SL partners) (involving 9 pairs, adjusted P=5.63E-04). These results suggest that the cellular immune responses induced by SARS-CoV-2 infection can potentially control the cells fate when targeted in conjunction with the central pathways regulating cell proliferation and nucleic acid/protein metabolism.

**Figure 2.**
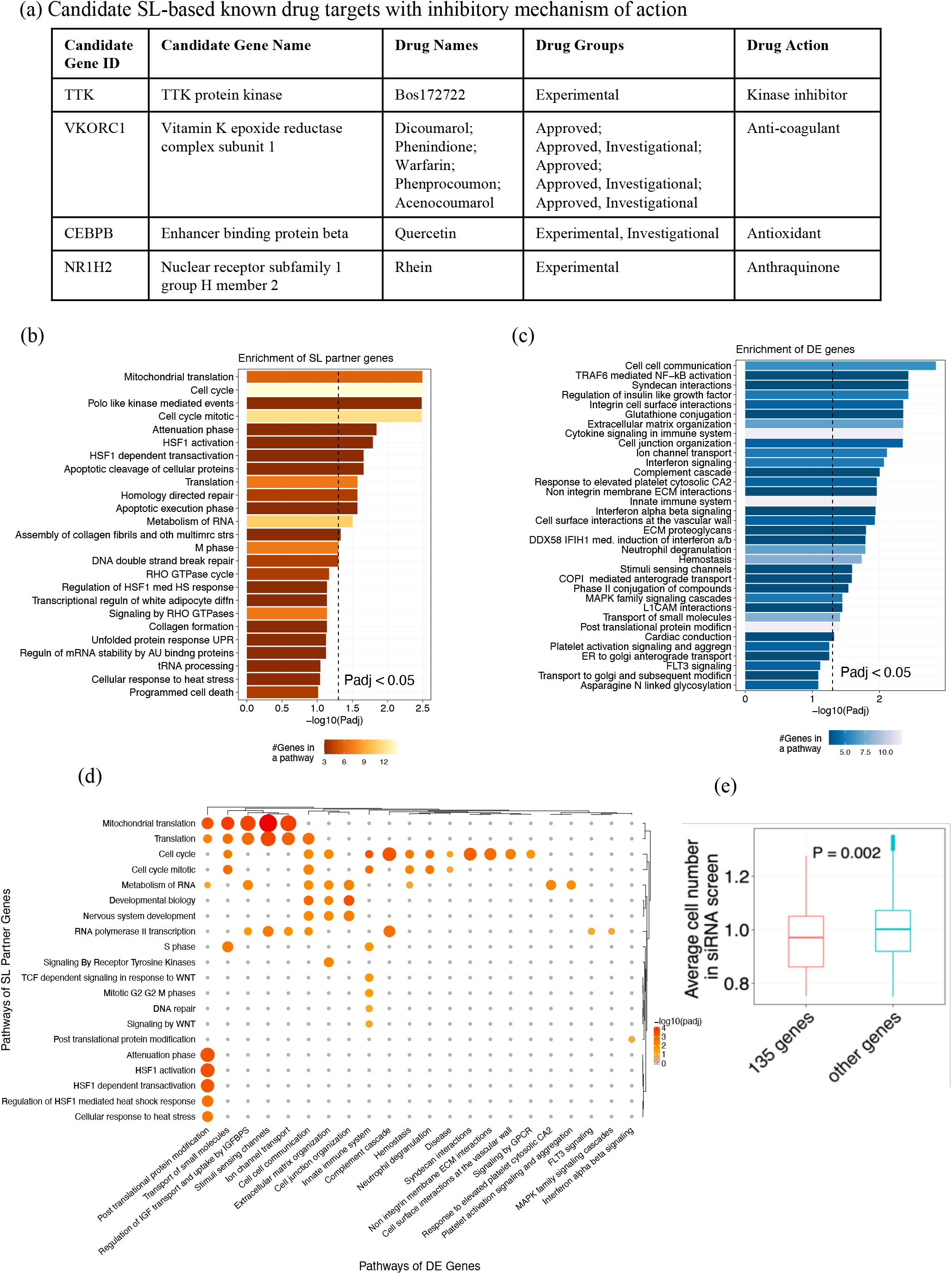
(a) A panel of candidate SL-based known drug targets with inhibitory mechanism of action. These drugs can be potentially repurposed to treat SARS-CoV-2 infection. **(b)** Pathway enrichment results for the 140 candidate SL-based targets. A bar plot showing the negative log10-transformed Benjamini-Hochberg-adjusted P values (Padj) from Fisher’s exact tests (X-axis) for the top pathways from the Reactome database (Fabregat et al. 2018) enriched by these SL-based targets. The color of the bars encodes the number of overlapping genes within each pathway. The black dotted line corresponds to the cutoff of adjusted P<0.05. **(c)** A similar plot as in (b) showing the top pathways enriched by the SARS-CoV-2-induced differentially expressed (DE) genes that form SL/SDL interactions with the 140 candidate target genes. **(d)** A heatmap illustrating the enrichment of DE gene - SL partner gene pairs formed by the 140 candidate targets in various combinations of pathways. Significant pathway combinations with adjusted P<0.1 are shown here. Pathways of the DE genes are given on the horizontal axis (columns), and pathways of the SL partner genes are given on the vertical axis (rows) of the heatmap. Size of the circles correspond to their odds ratio of enrichment and color of the circles correspond to the negative log10 adjusted P-values of a pathway combination. **(e)** Box plot of one-sided Wilcoxon rank-sum test results for average cell number between 135 candidate genes and the rest of the genes in the genome-wide siRNA screen. P-value is shown in between the boxes.

In the screens analyzed above, the cell viability-decreasing effects observed are potentially confounded by virus-induced cytopathic effects. We hence additionally analyzed a very recent imaging-based genome-wide siRNA screen that we performed previously in human Caco-2 cells (data deposited at https://figshare.com/s/4117ac39b1d21b56f5e6), where SARS-CoV-2 replication and cell number after each individual target gene knockdown were measured independently via anti-SARS-CoV-2 antibody and DAPI staining, respectively. 135 of our 140 predicted candidate genes were covered within the siRNA library of this screen (**Table S3H**). First, across these genes, we reassuringly found that knockdowns resulting in lower cell numbers tend to also have lower viral load (Spearman’s *ρ=*0.16, P<2.2e-16), supporting the notion that reducing the viability of the virus-infected cells can indeed inhibit virus replication. Second, we find that the knockdown of these 135 predicted genes indeed results in significantly lower cell numbers compared to the knockdown of the rest of the genes in the screen (one-sided Wilcoxon rank-sum test P=0.002, **Figure 2e**). Reassuringly, within the 135 SL candidates, the candidate genes with lower cell viability (relative cell number <1.0) have higher weighted DE score (one-sided Wilcoxon rank-sum test P=0.032), and similarly, the candidates with lower viral load (z score<0) also have higher weighted DE score (one-sided Wilcoxon rank-sum test P=0.057; Methods). These results suggest that the SL-based predictions and the associated weighted DE score measuring the strength of SL effects can facilitate the identification of anti-SARS-CoV-2 targets that function via specifically reducing the viability of the infected host cells.

### (D) Network identification and experimental testing of 26 top-ranked SL targets

Next, we aimed to further narrow down the list of targets that we should further test in an additional, dedicated genomic screen. To this end, we set to first identify the subset of SL interactions with higher relevance to SARS-CoV-2 infection. Using simulated annealing we identified an optimal SARS-CoV-2 specific SL subnetwork that maximizes the correlation between viral inhibition scores measured in the genome-wide siRNA screen conducted in house and the SL-based weighted DE scores of the target genes it includes (see Methods). To experimentally validate the effect of the resulting 26 candidates on SARS-CoV-2 replication and cell viability, we performed an additional imaging-based siRNA assay in Caco-2 cells (endogenous expression on *ACE2* and *TMPRSS2*, and the GI epithelium as a target for SARS-CoV-2). The individual knockdown of each of the 26 genes was performed in a 384-well plate format using four replicates per target (**Figure 3a**). The percentage of virus-infected cells after target knockdown (reflecting viral load) was quantified via immunofluorescence staining with anti-SARS-CoV-2 antibody. Non-targeting (scrambled) siRNAs were used as negative control, and siRNAs targeting *ACE2* and *TMPRSS2* (two genes essential for viral entry) as positive controls (**Figure 3b**; Methods). The total cell number after gene knockdown (reflecting cell viability) was measured based on DAPI staining, using siRNAs targeting essential genes for the survival of the host cells (Toxic siRNA) as positive controls for cell viability (**Figure 3c**).

**Figure 3.**
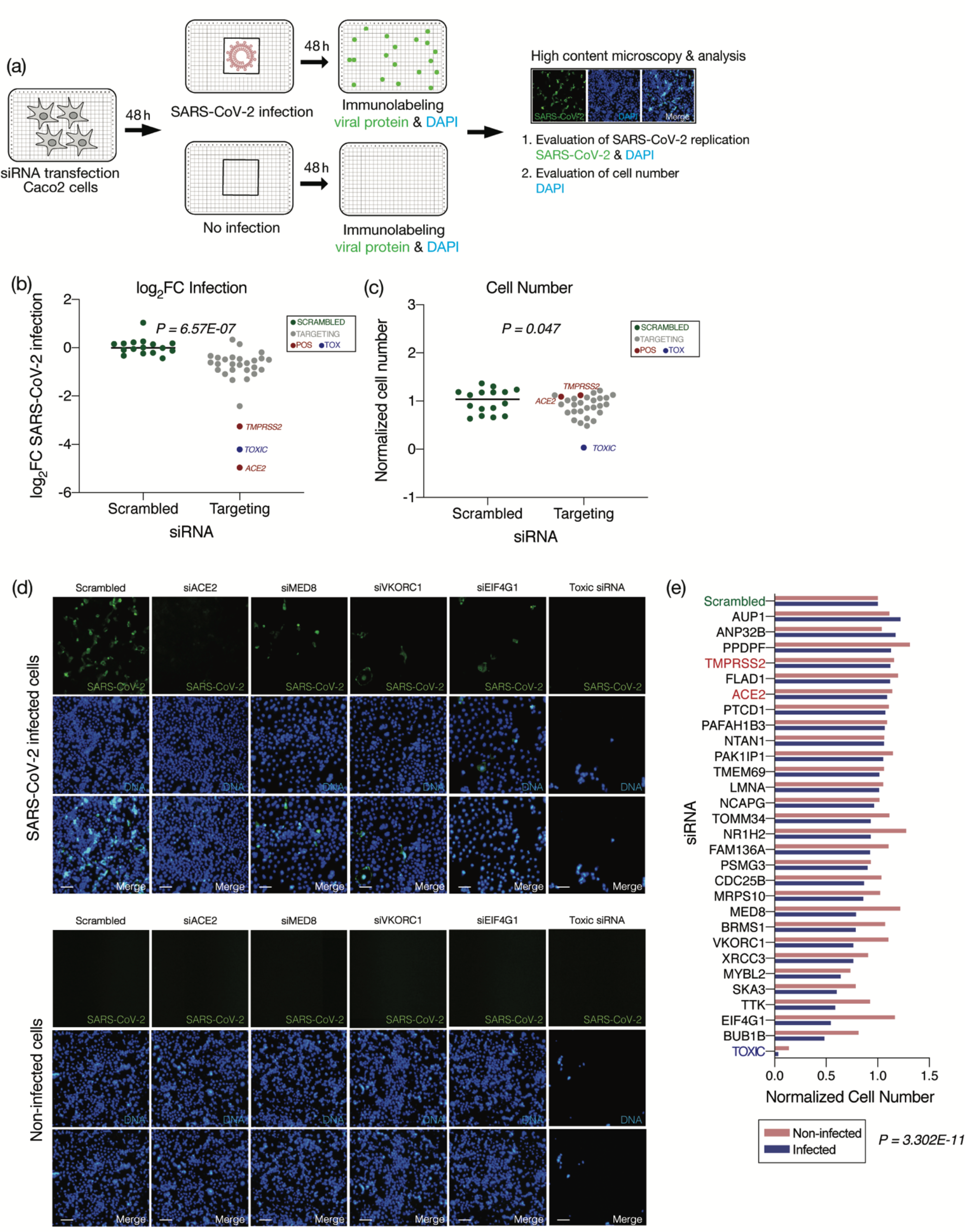
(a) Workflow of targeted siRNA screens in human Caco-2 cell with 4 replicates for each targeted gene. Cells were transfected with siRNAs and then either infected with SARS-CoV-2 (MOI = 0.1) or left non-infected. The number of cells (viability) after each gene knockdown was measured via DAPI staining (number of DAPI^+^ objects) and viral replication was evaluated via anti-SARS-CoV-2 antibody (percentage of SARS-CoV-2^+^ cells). (b) Scatter plot of viral infection in terms of average (n=4) log fold change of SARS-CoV-2 infected cells (log_2_FC infection values) of 26 SL targets (grey dots) relative to the scrambled siRNA (green dots). Positive controls ACE2 and TMPRSS2 are marked by red dots, and Toxic siRNA with blue dots. P-value from one-sided Wilcoxon rank-sum test between the scrambled siRNA vs. the rest of the genes is shown in the plot. (c) Scatter plot of cell viability in terms of average (n=4) normalized count of cell numbers from the DAPI-stained images of 26 SL targets relative to the scrambled siRNA. P-value from one-sided Wilcoxon rank-sum test between the scrambled siRNA vs. the rest of the genes is shown in the plot. (d) Representative images showing viral (SARS-CoV-2, green) and DAPI (DNA, blue) staining. Shown are cells treated with scrambled, siRNAs targeting positive controls (ACE2, TMPRSS2, and toxic siRNA), or 3 top target genes (VKORC1, MED8 and EIF4G1), and then either infected with SARS-CoV-2 (top panel) or left non-infected (bottom panel). Scale bar=10 μm. (e) Barplot of cell viabilities (average normalized cell numbers out of 4 replicates for each gene relative to scrambled) of siRNAs targeting each of the 26 SL targets and controls are plotted for both infected (blue) and non-infected (pink) conditions. One-sided Wilcoxon signed-rank test P-value of the average normalized cell numbers of each knocked-down target between two conditions is shown.

We first established the antiviral efficacy of the predicted targets - individual knockdown of the 26 predicted SL targets resulted in an overall reduction in viral replication (P=6.57E-07; **Figure 3b**) and reduced cell viability (P=0.047; **Figure 3c**) compared to scrambled siRNAs. Of note, the knockdown of *ACE2* and *TMPRSS2* also markedly reduced viral replication, consistent with their previously recognized potential as effective drug targets (Lin et al. 2021; Hoffman et al. 2020). Second, we tested if the predicted targets indeed selectively reduce the viability of the infected cells but not the non-infected cells. To this end we also measured cell number in siRNA transfected but non-infected cells (**Figure 3a**). Reassuringly, we found that the cell viabilities after knocking down the SL targets were significantly reduced in infected cells, but not in the non-infected cells (P=3.302E-11, **Figure 3d-e**). Overall, 24 out of 26 targets showed reduced viral load compared to the control and reduced cell viability in the infected condition compared to non-infected state (**Figure 3e**; **Table S4A**).

Among our 26 predicted targets, the top three emerging in our screen are *VKORC1*, an anti-coagulant that binds to ORF7a of SARS-CoV-2 (Holcomb et al. 2020; Janssen & Walk, 2020) and targeted by warfarin (Irwin et al. 2021); *MED8*, a gene required for the activation of transcription (Boube et al. 2002) and *EIF4G1*, known for its role in cell growth, proliferation and differentiation (Ramirez-Valle et al. 2008) (**Figure 3d**). Some of the validated targets are known to be involved in the context of other viral infections (**Table S4B**) – such as, *CDC25B* promotes influenza A virus replication (Cui et al. 2018).

## Discussion

Given the current lack of effective treatment options for COVID-19, identifying new drug targets for the virus causing this rapidly spreading disease with emerging new variants is highly critical. In this work, we predicted potential *clinically relevant* SL-based anti-SARS-CoV-2 targets. We further tested our initial list of 454 emerging predictions using two published CRISPR-Cas9 genetic screens in two different cell lines (Vero E6 and human A549 with exogenous *ACE2* expression) infected with SARS-CoV-2. Intersecting our predicted targets with the hits from the CRISPR screens, we further narrowed down this initial list to that of 140 clinically relevant candidate targets. A network-based analysis of these 140 targets using the results of a genome-wide siRNA screen that we have previously performed, led to the selection of a smaller subset of 26 genes, which we further experimentally tested and validated via a small-scale targeted siRNA screen in human Caco-2 cells, identifying targets that selectively reduce cell viability only in infected cells, demonstrating the utility of our synthetic lethality approach. Our analysis involves the integration of multiple published gene expression datasets of SARS-CoV-2 infection, both *in vitro* and *in vivo*, finding overall a moderate but robust level of concordance of virus-induced DE genes, especially and importantly across the *in vivo* datasets. This analysis encompasses some of the largest available transcriptomic datasets of COVID-19 patients (Butler et al. 2020, Lieberman et al. 2020), as well as single-cell datasets, which adds to the clinical relevance of our predicted targets. Many additional *in vitro* transcriptomics datasets on SARS-CoV-2 infection are available (like in human hepatoma cells), but we have chosen to build up our 140 targets list based on the two particular datasets of Vero E6 and A549^*ACE2*^ cells, as the same cell lines were used in the two CRISPR-Cas9 genetic screens that we used for validation.

Our study is motivated by the SL-based antiviral strategy proposed by Mast et al. (2020) with preliminary proof-of-concept application for poliovirus demonstrated in Navare et al. (2020). This strategy aims to specifically reduce the viability or impair the normal functioning of the infected host cells, thereby suppressing their capacity to support viral proliferation (Mast et al. 2020). To tease apart these two factors, we specifically employed an imaging-based siRNA assay where both viral replication and cell counts after gene knockdown were measured. In our previously reported genome-wide screen using this method, a positive correlation between these two variables was observed, supporting the inhibition of host cell viability as a potential antiviral strategy. In the targeted siRNA screen conducted in this study, a similar correlation was observed, where the knockdown of our 26 candidate targets significantly reduced viral replication accompanied by a reduction in cell number. However, importantly, knocking down these genes did not harm the non-infected cells. This reduction in cell viability in infected cells is likely not driven by virus-induced cytopathic effects (CPE), as cells treated with the negative control scrambled siRNA showed significantly higher rates of infection and higher cell number counts than cells treated with targeting siRNAs (**Figure 3b-d**). Further, the Pearson correlation value between log_2_FC infection and cell number in infected cells was in the positive range (*p value = 0*.*1235*). Obviously, further *in vivo* validations are required for the potential future therapy development based on these targets, specifically testing the potential toxicity issues that may arise with killing of infected cells. Even if such toxicity arises on a serious scale, SL-based treatments may possibly still prove useful at early stages of infection.

In summary, our study represents a first of its kind proof-of-concept application of a synthetic lethality inference approach to predict anti-SARS-CoV-2 targets in a systematic genome-wide manner. We provide a list of potential targets that are supported by multiple lines of evidence, importantly, emerging from the integrative analysis of both in vitro and in vivo data, giving rise to the hope that they then are more likely to be translationally relevant and help in the development of new drugs for treating COVID-19.

## Materials and Methods

### Differential expression analysis of transcriptomic data on SARS-CoV-2 infection

Several RNA-sequencing datasets of SARS-CoV-2-infected and matched control samples were obtained as follows: Vero E6 cells (Riva et al. 2020; from the GEO database, dataset ID GSE153940), A549 cells with exogenous *ACE2* (A549^*ACE2*^; Blanco-Melo et al. 2020; from GEO, GSE147507), nasopharyngeal swab samples from human COVID-19 patients (Butler et al. 2020 and Lieberman et al. 2020, from the Supplementary Materials of the respective publications), single-cell data of nasopharyngeal and bronchial samples from COVID-19 patients (Chua et al. 2020; from FigShare https://doi.org/10.6084/m9.figshare.12436517), and single-cell data of bronchoalveolar lavage fluid from COVID-19 patients (Liao et al. 2020; obtained following instructions in https://github.com/zhangzlab/covid_balf).

For the Vero E6 and A549^*ACE2*^ data, we performed differential expression (DE) analysis of virus-infected vs control samples with the DESeq2 R package (Love et al. 2014). Significant DE genes with FDR<0.05 were identified and further filtered by dataset-specific log fold-change cutoffs so as to get a reasonable number of such DE genes for further analysis. Specifically, 123 down (log fold-change< -1.0) and 171 up-regulated (log fold-change > 2.0) genes were identified from the Vero E6 dataset, 98 down (log fold-change< -1.5) and 172 up-regulated (log fold-change > 2.0) genes were identified from the A549^*ACE2*^ dataset. For the dataset from Butler et al. 2020, we used the virus-positive vs negative DE results provided by the authors that correspond to the analysis of 10 million human mapped reads with correction of potential confounders (batch effect and deconvolution of cell proportions). Similarly, we also used the DE results provided by the authors for the Lieberman et al. 2020 dataset. We combined the significant DE genes (FDR<0.05) with absolute log fold-change values greater than 1.0 from both datasets for further downstream analysis; this set includes 121 down and 166 up-regulated genes. For single-cell dataset from Chua et al. 2020 and Liao et al. 2020, we only focused on epithelial cells (based on the cell type annotation provided by the authors) as the virus-infected cells relevant to our target prediction approach and performed DE analysis for virus-positive vs negative samples using the Seurat R package (Stuart et al. 2019), specifically, for nasopharyngeal swab and bronchial basal cells from Chua et al. 2020 and bronchoalveolar lavage fluid epithelial cells from Liao et al. 2020. Significant DE genes (FDR<0.05) with absolute log fold-change values greater than 0.5 were selected, resulting in 166 down and 181 up-regulated genes from the Chua et al. dataset and 196 down and 320 up-regulated genes from the Liao et al. dataset.

### Identification of SL partners of the SARS-CoV-2-altered genes

For the significant DE genes identified from each of the above datasets, we identified their SL partners (for down-regulated genes) using the ISLE algorithm (Lee et al. 2018; **Table S2**). ISLE was originally designed for SL prediction in the cancer setting, but as discussed in the main text here we apply it to identify potential SLs for antivirus. ISLE involves four steps analyzing gene essentiality data in cancer cell lines, tumor molecular profiles, cancer patient survival data, and gene phylogeny data to identify clinically relevant SL interactions (described in detail in Lee et al. 2018). For the up-regulated DE genes, we instead identified their SDL partner genes also using ISLE but with minor modifications to account for the directional difference between SL and SDL. For example, in the third step of ISLE analyzing cancer patient survival data, while the original ISLE algorithm tests for the association of concomitant down-regulation of a pair of genes with better patient survival as a feature of SL, in the modification for SDL we instead tested for the association of down-regulation of one gene and the up-regulation of another gene in a gene pair with better patient survival, which corresponds to the SDL definition. We used FDR cutoff 0.1 to select significant SL/SDL partners. We ranked the ISLE-predicted partner genes by their P values from the patient survival analysis. Significant partner genes (a total of 454 genes, **Table S2F**) that are common to all analyzed datasets were taken for further downstream analysis.

### Prioritization of targets using genome-wide CRISPR-Cas9 genetic screening data in SARS-CoV-2-infected cells

We obtained two currently available genome-wide CRISPR-Cas9 genetic screening data aiming to identify host genes regulating SARS-CoV-2 infection, one from Vero E6 cell line (Wei et al. 2020) and the other from human alveolar basal epithelial carcinoma A549 cell line with exogenous *ACE2* expression (A549^*ACE2*^; Daniloski et al. 2020). The complete results of sgRNA differential expression were obtained from the Supplementary Materials of the two publications. To validate that our predicted targets are indeed SL with SARS-CoV-2 infection, i.e., the inhibition of these genes can lead to more cell dropout in the virus-infected cells than in control cells, we tested for enrichment of genes showing negative log fold-changes in the CRISPR-Cas9 screens with the GSEA method (Subramanian et al. 2005) as implemented in the R package fgsea (Korotkevich et al. 2019). Specifically, the log fold-change values across genes from each CRISPR-Cas9 screen were ranked, and GSEA was applied to the ranked list with the “gene set” being the 454 predicted SL-based targets common to all datasets. To obtain our final list of 140 candidate targets, we intersect the 454 consensus predictions with the genes showing the strongest negative log fold-changes in the CRISPR-Cas9 screens. These strongest CRISPR hits were selected as follows: In the Vero E6 dataset, we selected 680 genes with negative mean log2 fold-changes with FDR<0.1. The A549^*ACE2*^ dataset contains results from two screens with different multiplicities of infections (MOIs), and we selected 2811 genes from the lower MOI screen and 1653 genes from the higher MOI screen whose absolute log fold-changes are lesser than -0.5. The intersection of all selected CRISPR genes with the 454 consensus SL-based targets gives the final list of 140 candidate target genes (**Table S3**).

### Calculation of weighted DE score

Each of 140 candidates have the same or different significant DE genes from six different transcriptomic datasets - *in vitro* Vero E6 (Riva et al. 2020), *in vitro* A549^*ACE2*^ cells (Blanco-Melo et al. 2020), two *in vivo* Bulk Swab datasets (Butler et al. 2020 and Lieberman et al. 2020) and two *in vivo* single-cell RNA-seq datasets (Chua et al. 2020 and Liao et al. 2020). We hypothesized that the effect of knockdown of these SL-based targets is dependent upon the synthetic lethality strength, as reflected by the extent of perturbation of DE genes by the virus (log fold change) in their respective transcriptomic datasets. We considered the median absolute log fold change of a DE gene in all six datasets (if it is available). If a DE gene is down-regulated in the dataset where it is considered to be significant, then all down-regulated log fold changes in other datasets will contribute to the final median value, and similar will be the treatment for up-regulated cases. For a candidate gene g, its weighted DE score (S_g_) was calculated as in equation 1,

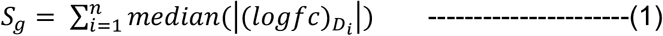

Where, ‘g’ is synthetic lethal partner of ‘n’ number of DE genes (D) and median log fold changes (in absolute value) of D_i_ from six different transcriptomic datasets were considered according to the status (either up-regulated or down-regulated) of the significant DE gene corresponding to the SL pair.

### Virus-interacting genes and pathway enrichment analysis on the predicted SL-based target genes

We also tested for the enrichment between the 454 predicted targets with human proteins reported to interact with SARS-CoV-2 viral proteins (Gordon et al. 2020 and Stukalov et al. 2020; data obtained from the Supplementary Materials of these publications) with Fisher’s exact test using the fisher.test function from the R software. Similarly, Fisher’s exact tests were used to test for pathway enrichment of the final SL-based candidate targets (**Table S3E**) and their corresponding DE genes (**Table S3F**) using pathway annotations from the Reactome database (Fabregat et al. 2018) collected from MSigDB v7.2 (Liberzon et al. 2011). Pathways with more than three genes overlapping with the candidate target set were considered. P values across pathways were corrected for multiple testing with the Benjamini-Hochberg method.

### Enrichment of pathway combinations for DE-S(D)L partner gene pairs

Fisher’s exact tests were used to test the enrichment of the SL gene pairs in each possible pathway combination (i.e. “pathway pair”) (**Table S3G**). Specifically, pathway annotation of single genes from the Reactome database (Fabregat et al. 2018) collected from MSigDB v7.2 (Liberzon et al. 2011) was used to create all possible pathway pairs and their member gene pairs. The “background” set for the enrichment was the gene pairs formed between the virus-induced DE genes and all genome-wide genes. P values were corrected with the Benjamini-Hochberg method.

### Annotation of the final SL-based candidate target gene list

The predicted targets (140 candidates) were mapped to known drugs (clinically approved and experimental/ investigational) that target (i.e. inhibit) them using drug target annotation from DrugBank (Wishart et al. 2018). We manually searched the literature on each of the 26 candidate genes to assess their relevance to SARS-CoV-2 or other viral infection based on prior knowledge. Detailed annotation with regard to literature support for the candidate genes are provided in **Table S4B**.

### Genome-wide siRNA screen

We used our previously performed genome-wide siRNA screen, carried out in human Caco-2 cells to identify host cell factors that affect the replication of SARS-CoV-2 (data deposited at https://figshare.com/s/4117ac39b1d21b56f5e6).

### Selection of 26 genes for experimental validation

Out of 140 candidate genes, we have cell viability and viral replication information for 135 SL-based targets. The size of our synthetic lethal network for 135 candidates including DE genes from all six different transcriptomic datasets was 854. We used the simulated annealing method to maximize the correlation between viral inhibition from our genome-wide siRNA screen and the SL-based weighted DE scores, which resulted in an optimal SARS-CoV-2 specific SL subnetwork. We finally selected 26 target genes from this subnetwork where viral load was reduced (z-score < 0) in the previously performed genome-wide siRNA screen.

‘Optim’ function in R was used to optimize the whole SL network, using variable network sizes and seed and using 10,000 iterations for each run. All other parameters were used as default in the function. Different sizes of networks were optimized for minimization of correlation (Spearman’s *ρ* in R) between weighted DE score and cell viability from siRNA screen for candidate genes present in that network. The resulting SL network was used to calculate the correlation between weighted DE score and viral load of those candidate genes in the selected network. The aim of this *in silico* experiment was to extract a SARS-CoV-2 specific SL network where cell viability and viral replication will be lower upon knocking down the candidate genes in that network.

### Targeted siRNA screen

#### Screen

To evaluate the effect of the 26 predicted targets on viral replication and cell viability, a targeted siRNA screen was conducted using human Caco-2 cells (ATCC). siRNAs targeting these genes were obtained from the Dharmacon ON-TARGETplus SMARTpool library and individually arrayed in 384-well plates at a concentration of 12.5 nM per well. First, siRNAs were incubated with 0.1 μL Lipofectamine RNAiMAX transfection reagent (ThermoFisher) diluted in 9.90 μl Opti-MEM media (ThermoFisher) for 20 min at room temperature. Then, 3,000 Caco-2 cells diluted in cell growth media (see below) were seeded on top and incubated for 48 h at 37°C with 5% CO_2_ conditions. Cells were then either infected with SARS-CoV-2 (isolate USA-WA1/2020) at a multiplicity of infection (MOI) 0.1 or left non-infected. After an incubation period of 48 h at 37°C with 5% CO_2_ conditions, cells were fixed with 4% PFA (Boston BioProducts) for 4 h at room temperature. Cells were then washed twice with PBS prior to 20 min permeabilization with 0.5% Triton X-100 and 1 h blocking with 3% BSA (Sigma) at room temperature. Cells were then incubated for 2 h at room temperature with primary anti-SARS-CoV-2 N protein rabbit polyclonal antibody (kind gift from Dr. Adolfo Garcia-Sastre), followed by three PBS washes and 1 h incubation with Alexa Fluor 488-conjugated anti-rabbit secondary antibody (A-11008, ThermoFisher) diluted in 3% BSA. Cells were washed three times with PBS and then stained with DAPI (4,6-diamidine-2-phenylindole, KPL). Plates were sealed and stored at 4°C until imaging. Caco-2 cells growth media: 40 μL DMEM media (Gibco) supplemented with 10% heat-inactivated fetal bovine serum (FBS, Gibco), and 50 U/mL penicillin - 50 µg/mL streptomycin (Fisher Scientific).

#### Analysis

The resulting number of cells and SARS-CoV-2 replication after each individual target knockdown was measured using high-content imaging. Images were acquired using the IC200 imaging system (Vala Sciences), and then analyzed using the analysis software Columbus v2.5 (Perkin Elmer). The number of DAPI^+^ objects was used to calculate the number of cells in the well, and the number of SARS-CoV-2^+^ to calculate the percentage of infected cells. The average percentage of infected cells for each target knockdown (n=4) was normalized to that of the negative control scrambled and used to calculate the log_2_FC infection.

## Acknowledgement

This research was supported in part by the Intramural Research Program of the National Institutes of Health, NCI, CCR; and used the computational resources of the NIH HPC Biowulf cluster (http://hpc.nih.gov). We acknowledge and thank the National Cancer Institute for providing financial and infrastructural support. The content of this publication does not necessarily reflect the views or policies of the Department of Health and Human Services, nor does mention of trade names, commercial products, or organizations imply endorsement by the U.S. Government. The following reagent was deposited by the Centers for Disease Control and Prevention and obtained through BEI Resources, NIAID, NIH: SARS-Related Coronavirus 2, Isolate USA-WA1/2020, NR-52281. This work was also supported by the following grants to the Sanford Burnham Prebys Medical Discovery Institute: DoD: W81XWH-20-1-0270; DHIPC: U19 AI118610; Fluomics/NOSI: U19 AI135972. K.C. and S.S is supported by the NCI-UMD Partnership for Integrative Cancer Research Program.

## Author contributions

LRP, KC, NUN, SS and ER conceived the study, collected the data, designed the computational analyses, performed the analyses and wrote the paper. LMS and SC designed the experimental analyses and wrote the paper. YP, LR, XY performed experimental analyses. FS and JSL helped in computational analyses. All authors approved the publication of this manuscript.

## Conflict of interest

The other authors declare no competing interests.

## Supplementary Tables

**Table S1**: Enriched Reactome pathways for DE gene sets (FDR<0.1).

**Table S2**: List of synthetic lethal pairs in each of the transcriptomic dataset used in this study and the list of unified 454 SL partner genes.

**Table S3**: List of 140 candidate targets and three subsets comprising known drug targets, hub SL partner genes when each one of them are paired up with multiple DE genes and genes with enriched Reactome pathways. Pathway enrichment analysis (FDR <0.1) for SL partner genes, DE genes and combination pathway enrichment analysis for DE gene-SL partner gene pairs for 140 candidates. List of 135 candidates overlapping with genome-wide siRNA screen.

**Table S4**: Targeted siRNA screen results for selected 26 targets and annotation of 26 candidate targets with reference to SARS-CoV-2 or any other virus in the literature.

## Notes

### Competing Interest Statement

The authors have declared no competing interest.

